# Geomicrobiology of small primary and secondary lava caves at Capulin Volcano National Monument, NM, USA

**DOI:** 10.64898/2026.11.27.769469

**Authors:** Joseph R. Hoberg, Anika Baloun, Jimmy Swift, Abigail R. Brown, Georgia N. Schneider, Patricia E. Seiser, Daniel S. Jones

**Author notes:** G. Schneider, Bureau of Land Management, Department of the Interior, Carlsbad, NM, USA.

## Abstract

Several planetary bodies in our solar system, including Mars and the moon, have diverse volcanic landscapes that are thought to contain lava tubes and other caves. Lava caves on Earth are important analogs for evaluating the habitability and potential for life in these basalt-hosted subsurface environments. However, most prior studies have focused on large lava tubes, and little is known about the geomicrobiology of the many smaller caves that also occur in volcanic landscapes. In this study, we characterized the nature and occurrence of caves and cave-associated microbial communities in Capulin Volcano National Monument (CAVO), northeastern New Mexico. While we did not find remnants of any lava tubes within the Monument, we characterized microbial communities in several small primary and secondary lava caves. High-throughput 16S rRNA gene and transcript sequencing showed that the most abundant taxa on the walls of the small caves include *Rubrobacter*, *Modestobacter*, *Deinobacterium*, *Crossiella*, Archaea in the family “Nitrososphaeraceae”, several cyanobacteria, and many populations that were unclassified at the genus and family level. Most of the abundant populations were present in both rRNA gene and rRNA transcript libraries, indicating that the majority of the cave-associated community members are active and growing rather than dead or dormant. Many of the microbial taxa in CAVO caves are known from larger lava tubes, although some more specialized lineages, such as those associated with trace gas oxidation in caves, were rare or absent. These findings indicate that small lava caves harbor widespread cave-associated microbial communities and may collectively represent an underappreciated habitat in volcanic landscapes.

## Introduction

Lava caves have significant geological, ecological, and historical importance. They provide evidence for the nature and history of volcanic eruptions, contain rare and beautiful mineral deposits, and often represent significant archaeological and cultural heritage sites (Calvari and Pinkerton 1999; Forti 2005; Riquelme et al. 2017; Calvari et al. 2024). Lava caves can serve as shelter for animals, including specialized and sometimes endemic species (Nitzu et al. 2018; Howarth and Stone 2020), and contain microbial ecosystems with unique communities and microbe-mineral interactions (Northup and Lavoie 2001; Miller et al. 2014; Prescott et al. 2022). For planetary science and astrobiology, lava tubes and other caves or cave-like features in Earth’s volcanic landscapes are valuable analogs for similar structures on Mars and other planetary bodies, which could contain clues about past or present extraterrestrial life and might someday provide shelter for human habitation (Léveillé and Datta 2010; Northup et al. 2011; Popa et al. 2012).

Caves are commonly defined as voids that are large enough to be navigable by humans (Mylroie 2019), and can form by diverse processes including dissolution of soluble bedrock, cooling lava, melting glaciers, erosional or tectonic processes, and other mechanisms (Palmer 2007). Lava caves are caves that form in volcanic rock, and are classified as either primary or secondary. Primary lava caves form by cooling and usually flowing lava, so that the void forms at approximately the same time as the surrounding rock (Kempe 2019). In contrast, secondary lava caves form by erosion or tectonic activity in older volcanic rocks (Kempe 2019). Lava tubes, sometimes known as pyroducts (meaning “fire conduit”), are the most important primary lava caves worldwide, and are often many kilometers in length (Lockwood and Hazlett 2010). Primary lava caves also include smaller caves of limited extent, such as lava molds, tumulus and blister caves, pressure ridges, and volcanic vents (Slettebak 1981; Walker 1991; Sam et al. 2020). Secondary lava caves include talus caves, tectonic fissures, pit craters, and erosional caves such as sea caves (Palmer 2007; Kempe 2019).

Previous work has shown that lava tubes contain distinctive microbial communities that are important for biogeochemical cycling, primary production, and mineral precipitation and dissolution (Northup et al. 2011; Popa et al. 2012; Gonzalez-Pimentel et al. 2021; Selensky et al. 2021). Numerous rRNA gene amplicon surveys of microbial mats and mineral deposits in lava tubes show recurrent groups across regions such as Actinobacteria (Actinomycetota), Alphaproteobacteria, and Acidobacteriota, and genera such as *Crossiella*, “wb1-P19”, *Nitrospira*, *Rubrobacter*, “Ca. Omnitrophus”, and others (e.g., Northup et al. 2011; Northup and Lavoie 2015; Gonzalez-Pimentel et al. 2018; Nicolosi et al. 2023; Medley et al. 2024; Devlin et al. 2025). Lava caves also contain novel microbial populations that have not been described (e.g., Northup and Lavoie 2015; Weng et al. 2022; Fishman et al. 2023). Microbial communities vary among and within caves, with assemblages shifting across microclimate and geochemical gradients (Prescott et al. 2022). Comparisons between cave communities and overlying soils show few similarities, suggesting environmental filtering and endemicity within caves (Lavoie et al. 2017). However, with few exceptions (e.g., Léveillé et al. 2000; Léveillé et al. 2007), most studies have explored the microbiology of large lava tubes but not small lava caves, so it is not currently known whether these same microorganisms and microbiological processes also occur in smaller basalt caves, and are therefore are more widespread in volcanic terrains beyond the large voids.

Capulin Volcano National Monument (CAVO) is an 800 acre (324 hectare) national monument in northeastern New Mexico that receives 80,00-100,000 visitors per year (Figure 1). The Monument is based around Capulin Volcano, a 54,200 year old cinder cone in the Raton-Clayton volcanic field (Zimmerer 2019) (Figure 1). Some small lava caves have been observed in the lava flows that emerge from the base on the cone, and larger caves have been reported in the surrounding region. However, there has not been a detailed cave inventory in the Monument (KellerLynn 2015), or any geobiological studies of lava caves in the area. We therefore performed a reconnaissance survey to explore microbial communities and geomicrobiological processes in lava caves in CAVO. We identified several small primary and secondary lava caves, and used culture-based analyses and 16S rRNA gene- and transcript sequencing in order to (i) characterize microbial communities in these small lava caves, (ii) compare how microbial communities differ by cave type and among wall and floor deposits, and (iii) evaluate how microbial communities from small primary and secondary caves at CAVO compare to microbial communities from larger lava tubes.

**Figure 1.**
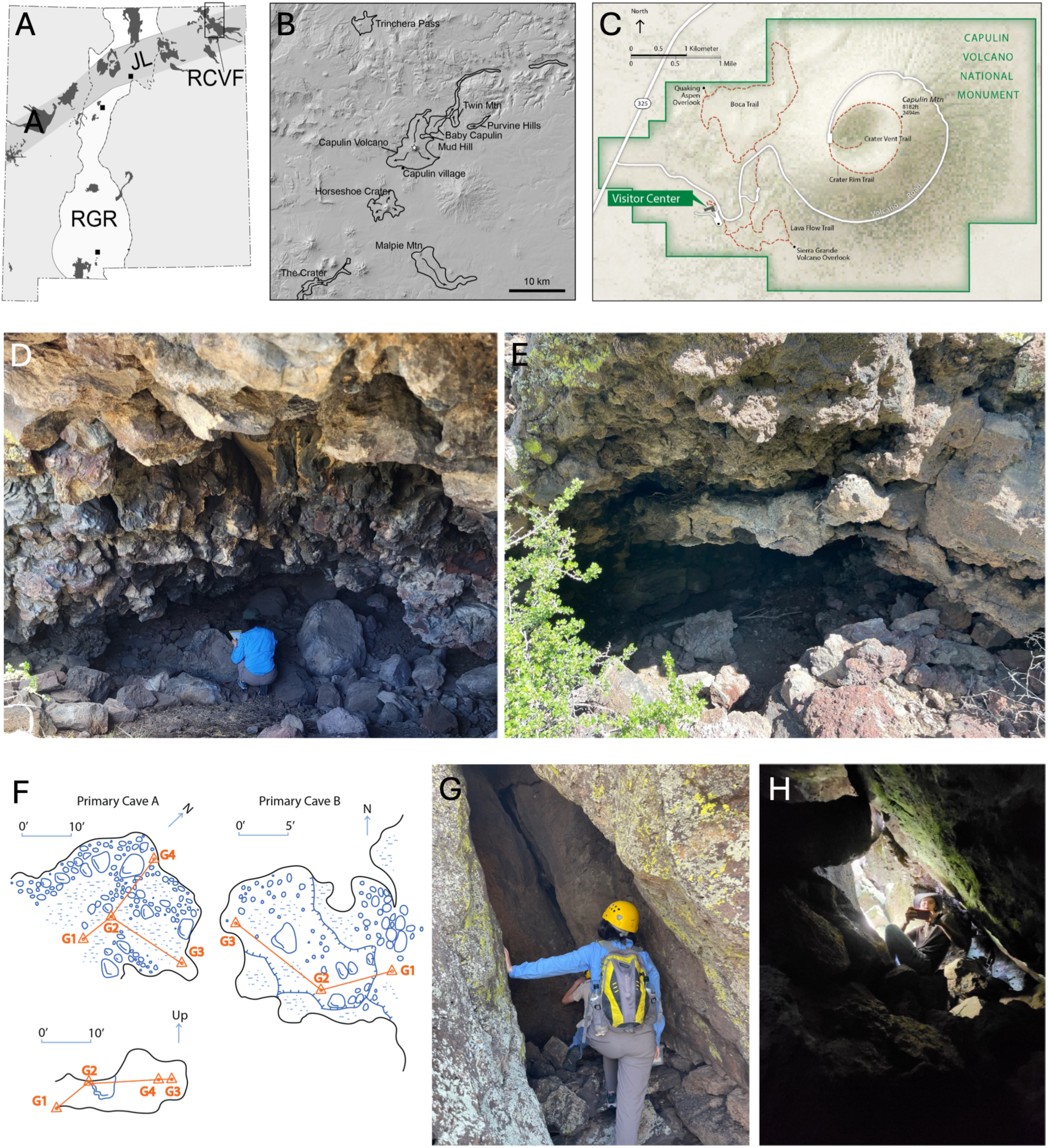
Geologic setting and small lava caves in Capulin Volcano National Monument. (A, B) Location of Capulin Volcano in the Raton-Clayton volcanic field (RCVF) (star in B); panel B is within the boxed area in A. RGR = Rio Grande Rift, JL = Jemez Lineament. Courtesy of M. Zimmerer. (C) Map of the Monument boundary (National Park Service). (D-F) Photos and maps of Primary Caves A and B. (G, H) Photos of Secondary Caves A and D. The people visible in the photos are authors Anika Baloun (photos A and G), Joseph Hoberg (photo G), and Jimmy Swift Jr. (photo H).

## Materials and Methods

### Site description and sample collection

Capulin Volcano National Monument (CAVO) is centered around a well-preserved cinder cone volcano that formed during the most recent phase of volcanic activity in the NM Raton-Clayton Volcanic Field approximately 54,200 ±1,800 years ago (Zimmerer 2019). The eastern, northern, and southern borders of the Monument end just outside the base of the cinder cone, but the western region includes volcanic terrain that was formed by lava flows that emerged from the base of the cone (Figure 1). There were four major flows that occurred during the eruptive episode, probably after the cinder cone itself was formed. These flows are known as the first, second, third, and fourth series, in order of occurrence: The earliest flows (first series) flowed to the east, then the second through fourth series flowed to the south, west, and north, in that order (Sayre and Ort 2011). According to Sayre and Ort (2011), the largest volume of lava came from the west side of the cone, and “formed the boca or mouth of the volcano, which includes several vents, tubes, lava lakes, lava cascades and levees”. This area, just to the north of the visitor center, includes hiking trails and picnic areas across the volcanic landscape (Figure 1). The composition of the lavas are basalt to basaltic trachyandesite. The cinder cone is mostly basaltic trachyandesite, while the basal flows are basaltic, with lower silicon, potassium, and incompatible trace elements compared to the trachyandesites (Ramos et al. 2019).

Samples were collected in June and July 2023 from cave floors or walls using sterilized metal scoopulas, spatulas, or tweezers to scoop or scrape sediment or mineral crusts into sterile tubes. Samples for nucleic acid extraction were immediately frozen and transported on dry ice, and then stored at −80°C in New Mexico Tech Geobiology Lab. Samples for enrichment culturing were stored in a cooler on water ice and then later at 4°C.

We did not find drip waters or pooled water inside any of the caves, so to characterize the chemistry of water that was in contact with the bedrock, water was sampled from small depressions on rock surfaces. pH, conductivity, temperature, and oxidation-reduction potential were measured on site with a YSI ProPlus multimeter (Xylem Inc., Washington DC, USA), and dissolved oxygen was measured with a YSI ProSolo optical dissolved oxygen probe (Xylem Inc.). Measurements of photosynthetically active radiation (PAR) inside caves were collected with a LI-250Q light meter (LI-COR Inc., Lincoln, NE, USA) in April 2026.

### Enrichment culturing and isolation

Sediments were enriched and isolated on solid R2A media (BD Difco™, Franklin Lakes, NJ). Enrichments were grown at room temperature. Colonies on solid media were plated at least twice to ensure isolation, and cultures were identified using direct PCR with primers 27F and 1492R as in Havlena et al. (2024). Successful gene products were cleaned with the DNA Clean & Concentrator kit (Zymo Research Corp., Irvine, CA, USA), and end-sequenced at GENEWIZ (Azenta Life Sciences, Burlington, MA, USA). Enrichment of ammonia oxidizers and anaerobic iron oxidizers were performed using media modified from Popa et al. (2012) and 1583 medium (DSMZ), and enrichment for aerobic iron oxidizers was performed using gradient tube techniques as in Hobart et al. (2025).

### Molecular methods

DNA extraction was performed as in Havlena et al. (2024), using the DNeasy PowerSoil Pro Kit (Qiagen, Hilden, Germany) with bead beating to ensure adequate DNA recovery. Samples were bead beat for 20, 40, and 60 s at 2500 RPM with a Mini Beadbeater (Biospec Products, Bartlesville, OK, USA), and then recombined to reduce DNA extraction bias. rRNA gene libraries were prepared by amplifying the V4 region of the 16S rRNA gene as in Jones et al. (2017). Most libraries were prepared using the ‘full service’ method with primers 515f and 806r, with some prepared using the ‘in house’ process and primers “515f modified” and 806r (Walters et al. 2015) with 30 or 35 PCR cycles as in Jones et al. (2017). DNA extraction blanks were checked for contamination using 35 PCR cycles, and re-extracted if any contamination was detected.

RNA was extracted as in Jones et al. (2017) and Best et al. (2025) using the RNeasy PowerBiofilm Kit (Qiagen) with 20, 40, and 60 s of beadbeating as for DNA extraction, above. DNA was removed from using the on-column treatment with DNase I as part of the RNeasy kit, and then a second DNase treatment using the TURBO DNA-free Kit (Invitrogen, Thermo Fisher Scientific, Waltham, MA, USA) and then cleaned with the RNA Clean and Concentrator-5 Kit (Zymo Research, Irvine, CA, USA). rRNA transcripts were amplified with the OneStep Ahead RT-PCR Kit (Qiagen) using primers 515f modified and 806r and either 25 or 30 cycles. To check for residual DNA, the RNA extract was amplified for 35 PCR cycles as above using the same V4 primers; if DNA amplification was observed, a third DNase treatment was performed with the TURBO DNA-free Kit as described above.

rRNA gene and transcript libraries were sequenced at the University of Minnesota Genomics Center on an Illumina MiSeq (2×300 paired end cycles, version 3 chemistry). ‘Full service’ libraries were created from DNA extracts as in Gohl et al. (2016), and PCR products for ‘in house’ libraries (which included some rRNA gene libraries and all rRNA transcript libraries) were barcoded and pooled for sequencing with the other libraries. Despite having no visible PCR product, DNA and RNA extraction blanks and PCR blanks were submitted for sequencing with sample libraries, along with some of the positive controls from PCR reactions.

### Statistical and bioinformatic analyses

Bioinformatic and statistical analyses, including operational taxonomic unit (OTU) calling and multivariate analyses, were performed as in Havlena et al. (2024). Raw sequences were trimmed to quality ≥ 28 and length ≥100 using Sickle (https://github.com/najoshi/sickle), any remaining adapters at the 3′ end were cropped with cutadapt (Martin 2011), forward and reverse reads merged with PEAR (Zhang et al. 2014), and operational taxonomic units (OTUs) were called at 97% similarity with chimera removal using USEARCH v11 (Edgar 2013) and VSEARCH v2.21 (Rognes et al. 2016). OTUs were classified with mother v1.36.1 (Schloss et al. 2009), using the Silva v138 database (Pruesse et al. 2007) and a confidence score cutoff of 50.

In order to compare rRNA gene libraries from this study to rRNA gene libraries from larger lava tubes, we selected rRNA gene libraries from three studies that used similar primers sets to characterize communities in caves from three different geographic areas, and that had publicly accessible rRNA gene libraries. Weng et al. (2022) and Fishman et al. (2023), which characterized lava tube communities from Craters of the Moon National Monument, ID, USA, and Mauna Loa, HI, USA, respectively, used the same V4 primer sets as we used here although produced shorter reads, while O’Connor et al. (2021) used primer sets 515f and 926r that cover the V4 and V5 region (Parada et al. 2016) to characterize microbial communities from Lava Beds National Monument, CA, USA. Therefore, compare among the four studies, we only used the first 125 bp of the R1 (forward) read, which encompasses part of the V4 region of the 16S rRNA gene. For CAVO samples, only rRNA gene libraries were used in this comparison. Quality filtering and trimming, OTU calling, and classification was otherwise performed as described above.

Statistical analyses were performed in R (R Core Team 2025) with the vegan package (Oksanen et al. 2025). Prior to analysis, libraries with fewer than 10,000 reads were removed. We also removed four libraries that were similar to blanks and were therefore assumed to have been impacted by contamination (Table S1). Raw OTU counts were first converted to proportions, rare OTUs (<0.01% of the total) were removed, and then proportional values were transformed using the Hellinger transformation, *b_ij_ = x_ij_^-0.5^*, where *x_ij_* is the proportion of object *x* in the OTU matrix and *b_ij_* is transformed object. Nonmetric multidimensional scaling (NMDS) ordinations were performed using the metaMDS() function with three dimensions (k = 3) for CAVO libraries only and four dimensions (k = 4) for the comparison to other lava tubes. Hierarchical agglomerative cluster analyses were calculated with Euclidean distance and Ward’s linkage using the functions *vegdist* and *agnes* functions in the vegan and cluster packages (Maechler et al. 2025). Q-mode cluster analysis (clustering of samples) was calculated for all OTUs, and R-mode analysis (clustering of OTUs) was performed with the top 50 or 75 most abundant OTUs. ANOSIM and PERMANOVA were performed with the vegan package using default parameters with the *anosim* and *adonis2* functions, with *pairwise.adonis* for pairwise comparisons (Martinez Arbizu 2020).

Raw sequences are available on the National Center for Biotechnology Information (NCBI) Sequence Read Archive (SRA) under BioProject PRJNA1567245.

## Results

### Cave descriptions and field observations

We collected samples from seven small lava caves in the monument. While there is no universally-accepted definition of a cave (Mylroie 2019), the most widely-applied definition is that a cave is a void navigable by a human. Therefore, for the purposes of this study, we used an inclusive criterion and defined a cave as a void large enough to easily fit our smallest group member, at least roughly 2 m deep by 0.5 m wide and 0.5 tall. None of the caves that we sampled here have fully dark zones that are accessible to humans, and light readings ranged from 5.93 to 0.01 μmol photons m^-2^ s^-1^ (Table S2), similar to the twilight zone (and lower than the entrance zone) of larger caves (e.g., Monro et al. 2018). While we use the term ‘cave’ here, we note that some of these structures might be considered ‘shelters’ or ‘shelter caves’.

We mapped two primary lava caves, which we refer to as Primary Cave 1 and Primary Cave 2. Both are small dome-shaped caves, up to several meters wide and less than two meters high (Figure 1d-f). We also observed several secondary lava caves, mostly talus caves within the lava flows, which in some places have many large boulders that create substantial subsurface void space, although not that is always accessible to humans. In total, we sampled in five secondary lava caves. Four of the secondary caves did not have a noticeable temperature difference from shaded outside areas, but one cave, Secondary Cave D, was cool and moist at depth (Figure 1g,h). We also note that, while there are features described as collapsed lava tubes on geologic maps (Sayre and Ort 2011; KellerLynn 2015), we did not discover any evidence of voids on the margins, upstream, or downstream edges of these features as would be expected for collapsed lava tubes.

We collected samples from the floor and ceilings of the two primary and five secondary caves. The floor sediments were loose, dusty, buff-colored, and fine-grained, and were easily scooped into sampling tubes. The cave walls sometimes contained green films, dark brown sediments, and thin white mineral crusts, sometimes dry and sometimes damp (Figure 2). There was evidence of animal use: many caves had packrat droppings, the largest primary cave had possible packrat urine, and evidence for bat use from guano and moth wings were noted in one of the secondary caves (B. Tobin, *unpublished observations*).

**Figure 2.**
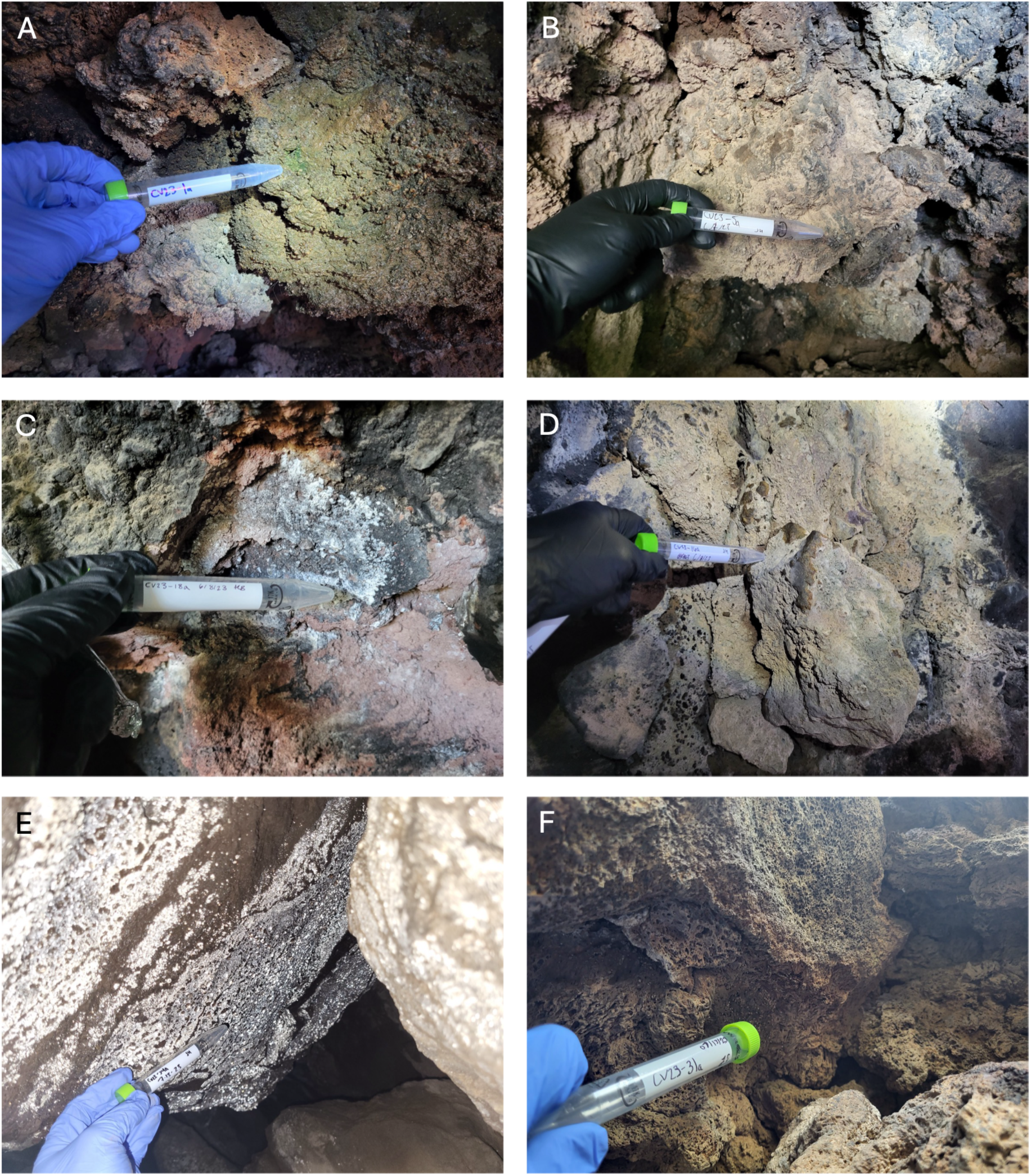
Photos of cave wall crusts and other wall features sampled. (A, B) Green wall sediments and buff-colored wall crusts in Primary Cave B; (C, D) white and brown wall crusts in Primary Cave A; (E, F) white and brown wall crusts in Secondary Caves D and C, respectively.

There were precipitation events around the time that samples were collected, and we found small pools on rock surfaces near two of the caves that we sampled to characterize the chemistry of water that had been in contact with the volcanic rocks. A pool near Primary Cave B had a pH value of 6.61, specific conductivity of 58.9 μS cm^-1^, oxidation reduction potential (ORP) of 67.3 mV, and temperature of 23.9°C at the time of sampling. In contrast, a pool near Secondary Cave B had a pH value of 9.41, specific conductivity of 51.8 μS cm^-1^, ORP of 39.7 mV, dissolved oxygen 11.8 mg L^-1^, and temperature of 23.7°C at the time of sampling. This pool contained a visible green film.

### Enrichment culturing and isolation

We enriched and isolated 18 bacterial strains on R2A media. Several were most closely related to *Bacillus* spp. or its closely related genera *Neobacillus* and *Paenibacillus*, and others had top BLASTN matches to *Arthrobacter*, *Pontibacter*, *Curtobacterium*, *Pseudomonas*, *Massilia*, *Dietzia*, *Burkholderia*, *Blastococcus*, and *Streptomyces* (Table S3). We also attempted to enrich autotrophic ammonia and iron oxidizers. Growth was inconclusive in media for aerobic ammonia oxidizers and anaerobic iron oxidizers, but a clear growth band was observed in gradient tubes for iron oxidizers. However, enrichment efforts for lithoautotrophs were abandoned before the enriched microorganisms were identified.

### 16S rRNA gene and transcript libraries

We generated 36 successful rRNA gene libraries from the two primary and five secondary caves, which had between 21,046 and 156,291 reads (mean reads 88,620, standard deviation 42,195) after quality trimming and filtering. We also generated 21 successful rRNA transcript libraries, which had between 25,270 and 65,202 reads (mean 42,894, standard deviation 11,523).

The most abundant OTUs include several Actinobacteriota classified as the genera *Rubrobacter*, *Modestobacter*, *Crossiella*, *Blastococcus*, *Arthrobacter,* and *Actinomycetospora*, as well as OTUs classified as members of the Proteobacteria and Deinococcota such as *Bradyrhizobium* spp., *Acidiphilium* spp., and *Deinobacteria* spp. (Figure 3). Many OTUs from these phyla were unclassified at the genus and family levels (Figure 3). There were also several abundant Archaeal OTUs classified as *Halococcus* spp. and an unclassified member of family Nitrososphaeraceae. The libraries also included abundant mitochondria and chloroplasts, as well as OTUs classified as non-chloroplast Cyanobacteria such as *Scytonema* and unclassified members of family Chroococcidiopsaceae.

**Figure 3.**
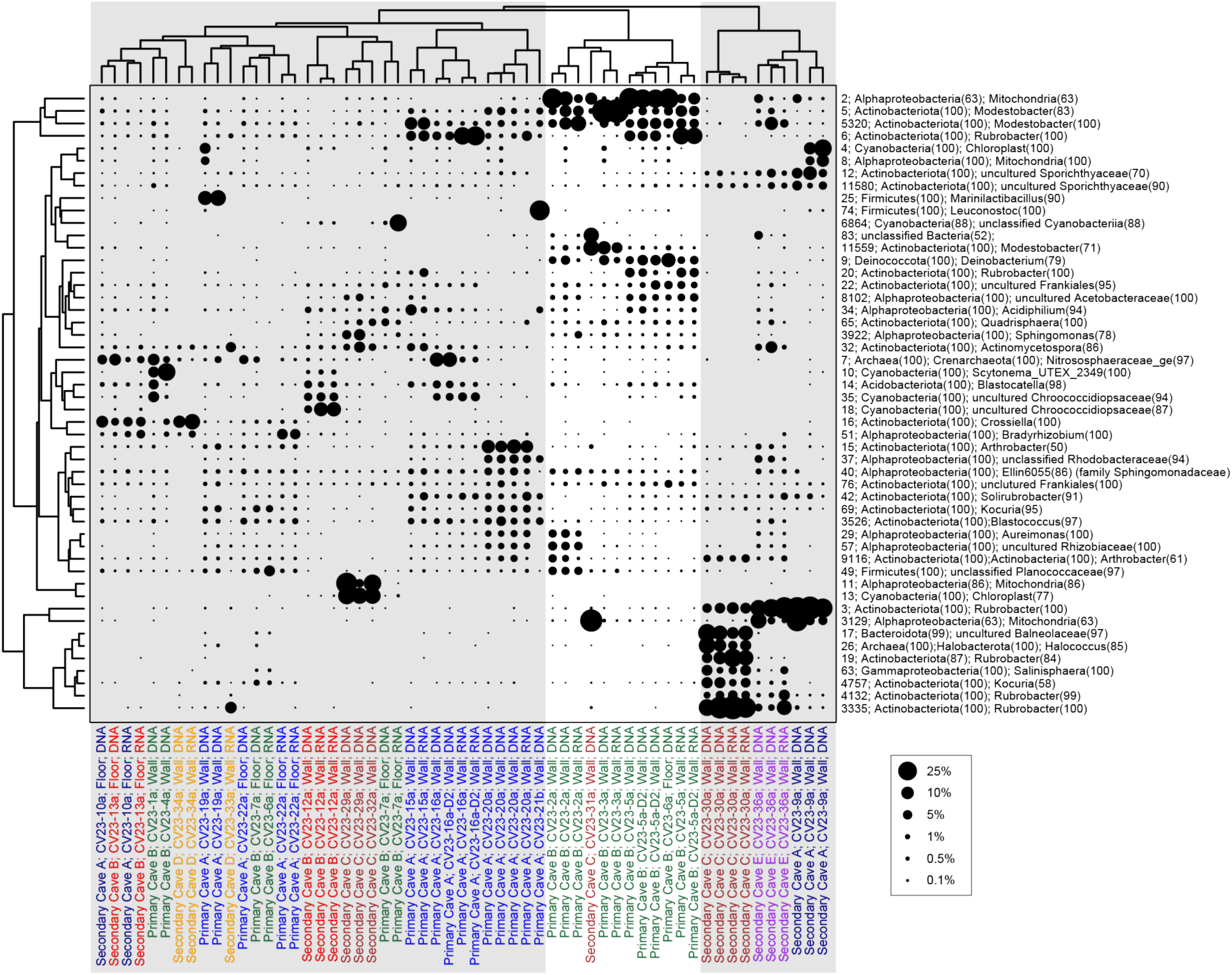
Two-way hierarchical agglomerative cluster analysis of 16S rRNA gene and transcript libraries. The sizes of the points scale with the relative abundance of the OTUs. The Q-mode cluster analysis was calculated with all OTUs, and sample labels are color coded by cave. R-mode clustering was calculated with the 50 most abundant OTUs.

These OTUs are present in both rRNA gene and rRNA transcript libraries (Figure 3), indicating that these populations are not only present but alive and active in most samples. Indeed, rRNA transcript libraries are overall similar to rRNA gene libraries, often most rRNA gene and transcript libraries from the same sample clustering together in hierarchical agglomerative cluster analysis and non-metric multidimensional scaling (NMDS) ordinations (Figures 3 and 4).

**Figure 4.**
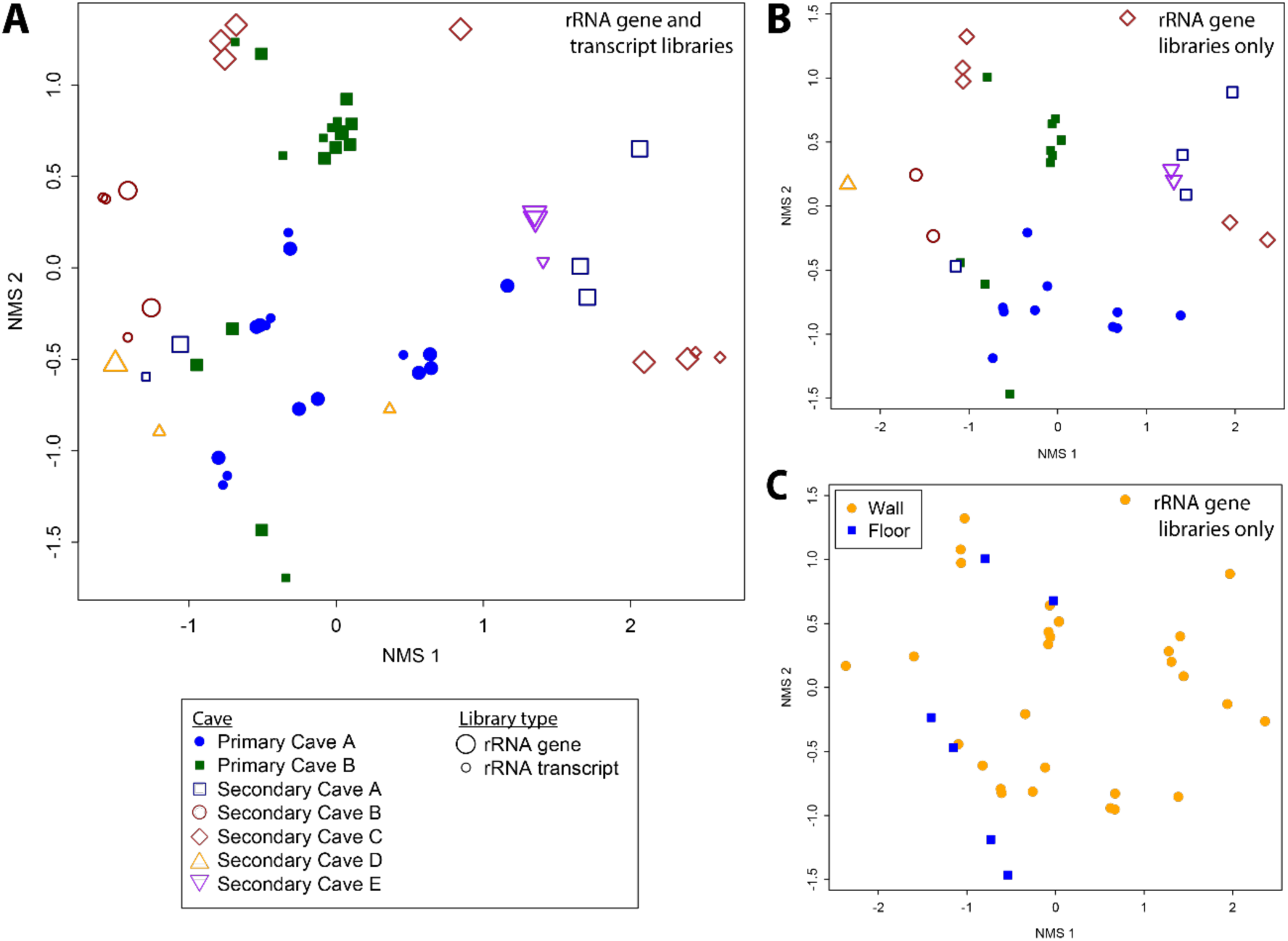
Non-metric multidimensional scaling (NMS) ordinations. Panel A includes rRNA gene and transcript libraries from CAVO, with larger points indicating rRNA gene libraries and smaller points indicating rRNA transcript libraries. Panels B and C show rRNA gene libraries alone, with points colored by cave (B) and wall deposits versus floor sediments (C). Stress is 6.4 and 7.3 for the ordinations in A and B/C, respectively.

Overall, rRNA gene and transcript libraries from primary lava caves were statistically significantly different from those in secondary caves (ANOSIM R = 0.337, p < 0.001; PERMANOVA F = 5.22, p < 0.001), and that remains true if only rRNA gene libraries or only rRNA transcript libraries are compared (rRNA gene libraries: ANOSIM R = 0.323, p < 0.001; PERMANOVA: F = 3.45, p < 0.001; rRNA transcript libraries: ANOSIM R = 0.306, p = 0.002; PERMANOVA: F = 2.63, p < 0.001). NMDS ordination shows that samples from the two primary caves cluster in the center of the ordination, while libraries from the secondary caves occur at the extremes of the first ordination axis; this pattern occurs with both rRNA gene and transcript libraries (Figure 4a) as well as with rRNA gene libraries alone (Figure 4b). rRNA gene libraries of sediments from the cave floors cluster with samples from cave walls (Figure 4c), and are not statistically significantly different (ANOSIM R < 0.1, p > 0.05 for rRNA gene libraries or both rRNA gene and transcript libraries).

### Comparison of CAVO cave communities to those from large lava tubes

We compared 16S rRNA gene libraries from the small primary and secondary lava caves at CAVO to publicly accessible libraries from three studies of large lava tubes: Lava Beds National Monument, CA, USA (O’Connor et al. 2021), Craters of the Moon National Monument, ID, USA (Weng et al. 2022), and Mauna Loa, HI, USA (Fishman et al. 2023). These comparisons were performed with the first 125 bp of forward reads only. Non-metric multidimensional scaling (NMDS) ordination shows that rRNA gene libraries from CAVO separate from those from the three lava tubes along the first and second ordination axes (Figure 5). However, some of the libraries from the primary caves are closer in ordination space to the lava tube libraries than libraries from the secondary caves.

**Figure 5.**
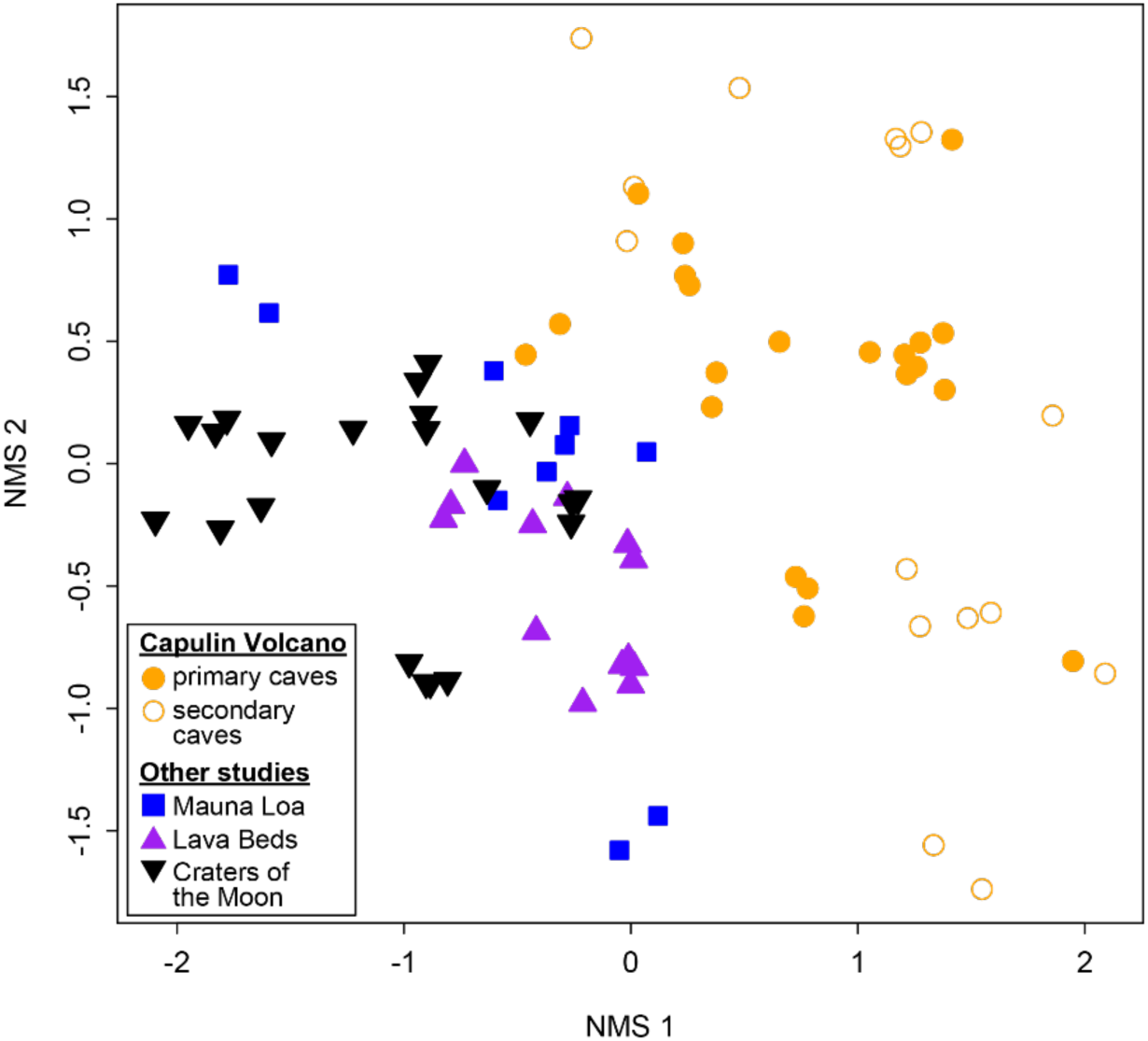
Non-metric multidimensional scaling (NMS) ordination comparing rRNA gene libraries from Capulin Volcano to rRNA gene libraries from large lava tubes from three other studies: Mauna Loa, HI, USA (Fishman et al. 2023), Lava Beds National Monument, CA, USA (O’Connor et al. 2021), and Craters of the Moon, ID, USA (Weng et al. 2022). Stress = 8.3.

To further explore similarities and differences among communities from the large lava tubes to small CAVO caves, we used two-way cluster analysis to explore how the most abundant taxa are distributed among the samples. In the R-mode analysis (clustering of taxa), clusters labeled I, V, and IX represent assemblages of OTUs that were most abundant in CAVO samples but were still present in the other caves. Clusters I contains OTUs with high confidence classifications to *Rubrobacter*, *Blastocatella*, and *Actinomycetospora*, and lower confidence assignments to the genera *Blastococcus*, *Deinobacterium*, *Acidiphilium*, *Geodermatophilus*, and *Modestobacter*, as well as some OTUs that are unclassified at the genus level. Cluster V contains OTUs with lower confidence assignments to *Paenisporosarcina*, *Aureimonas*, *Kocuria*, *Blastococcus*, and unclassified members of family Rhodobacteraceae and Micrococcaceae. Cluster IX contains OTUs classified as *Rubrobacter*, unknown Sporichthyaceae, and mitochondria.

In contrast, Cluster VIII contain OTUs that are found in the other caves but are still present at low abundance at CAVO, while Clusters IV and VII contain OTUs that are present in the other caves but are absent or rare in CAVO libraries. Cluster VIII contains OTUs with high confidence assignments to the genera *Rubrobacter* and “AKYG1722_ge” (phylum *Chloroflexiota*), low confidence assignments to the genera *Conexibacter*, *Quadrisphaera*, *Crossiella*, “JG30-KF-CM45_ge” (phylum Chloroflexiota), *Aliihoeflea*, and *Brevundimonas*, as well as several OTUs that are unclassified at the genus or family levels. Cluster IV contains OTUs classified as *Flavobacterium*, *Rhodococcus*, *Algoriphagus*, and unclassified Flavobacteriaceae, while Cluster XII contains “wb1-P19” and unclassified members of families Nitrosococcaceae and Beijerinckiaceae in the Proteobacteria.

## Discussion

### Occurrence and nature of caves in Capulin Volcano National Monument

We identified two primary and five secondary caves within the Monument (Table 1, Figure 1). The primary lava caves are most likely tumulus or possibly gas blister caves, formed by injection of lava or gases, respectively, from underneath a crusted lava surface (Walker 1991; Grimes 2002; Gadányi 2010). The former may be more likely, because the two caves have relatively thick roofs, roughly one meter, but they also have a dome-like structure more typical of blisters (Figure 1). They also could have formed by lateral compression of the crust over a flow, which can form similarly small lava caves by buckling of the crust (Grimes 2002). It is worth noting that there is substantial confusion of terminology used for these smaller lava caves, with terms such as stony rises, pressure ridge caves, lava blisters, gas blisters also present in the literature (e.g., Walker 1991; Grimes 2002; Palmer 2007).

**Table 1.**
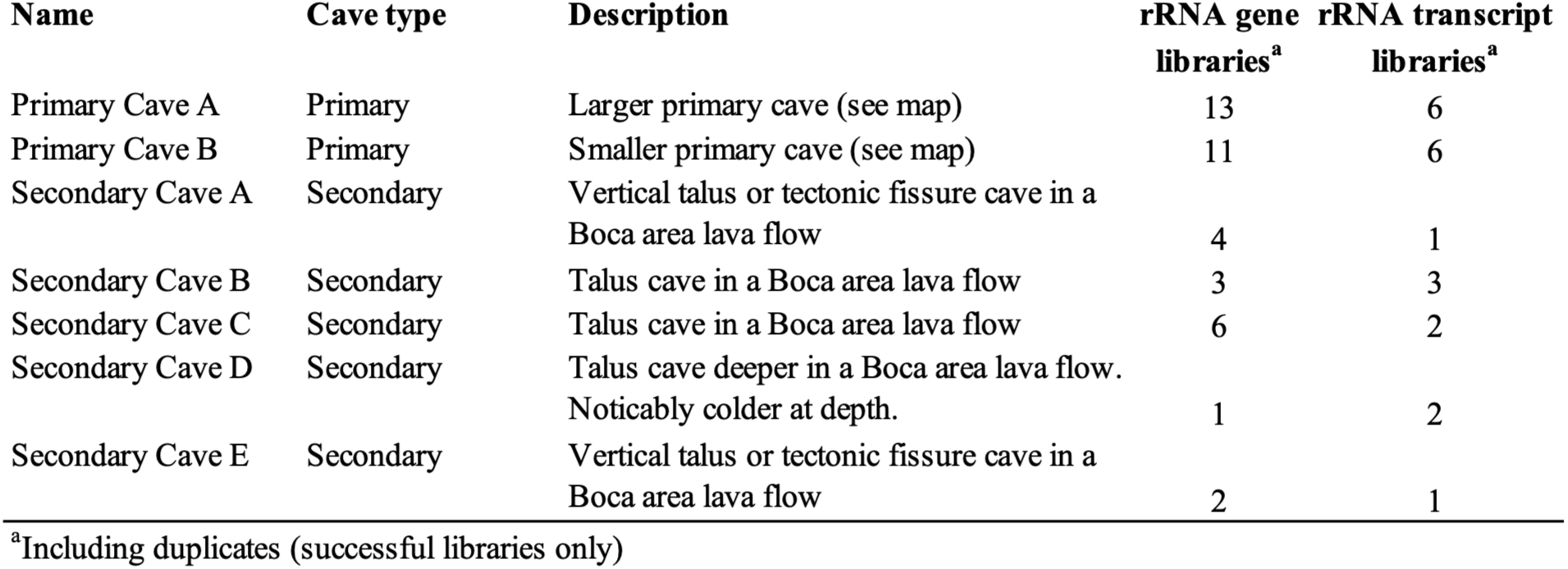
Summary of caves sampled.

Secondary caves were found within the lava flows that emerge from the western edge of the cinder cone. The largest cave, Secondary Cave D, was noticeably cooler at depth than other caves, and the cold and moist environment in this cave may be more similar to that of larger lava caves (although see the discussion on microbial communities, below). It is likely that there are similar and even more extensive caves within these boulder fields, as well as substantial subsurface void space that is not directly accessible by humans but that could represent subsurface access and shelter for subterranean-adapted fauna, and may contain cave-like microbial communities. Indeed, evidence of bat and packrat use was noted, and other studies have identified troglobitic invertebrates in large talus fields (Cokendolpher and Krejca 2010).

### Total and active microbial communities in small primary and secondary lava caves

The small lava caves sampled here have diverse microbial communities. While there is substantial variance among individual samples, there does not appear to be any clear trends among individual caves or between primary and secondary caves, and surprisingly, there are not clear difference between microbial communities from floor sediments versus wall deposits (Figure 4). Two-way cluster analysis shows three major clusters among the most abundant taxa, with more than half the samples occurring in one large cluster that has relatively consistent proportions of the most abundant OTUs (Figure 3).

Most of the OTUs occur in both rRNA gene and rRNA transcript libraries, which shows that most microbial populations that are present are also alive and active. This is in contrast with environments that have many inactive populations, which appear in rRNA gene but not rRNA transcript libraries, such as certain sedimentary environments or municipal drinking water (e.g., Revetta et al. 2011; Kearns et al. 2016; Li R et al. 2017; Reese et al. 2018). This may indicate that there is enough energy influx into the small lava caves to support robust microbial communities. For example, the presence of cyanobacteria and chloroplasts in many samples (Figures 3 and 7) is consistent with greater light influence in these small caves, and the transport of organic materials is likely also higher than would be typical for most deep cave environments. Furthermore, this finding suggests that the abundant microbial populations in these small lava caves are not simply allochthonous cells blown in from the surface that are persisting in an inactive state, but are instead active and established in the shallow cave environment.

**Figure 6.**
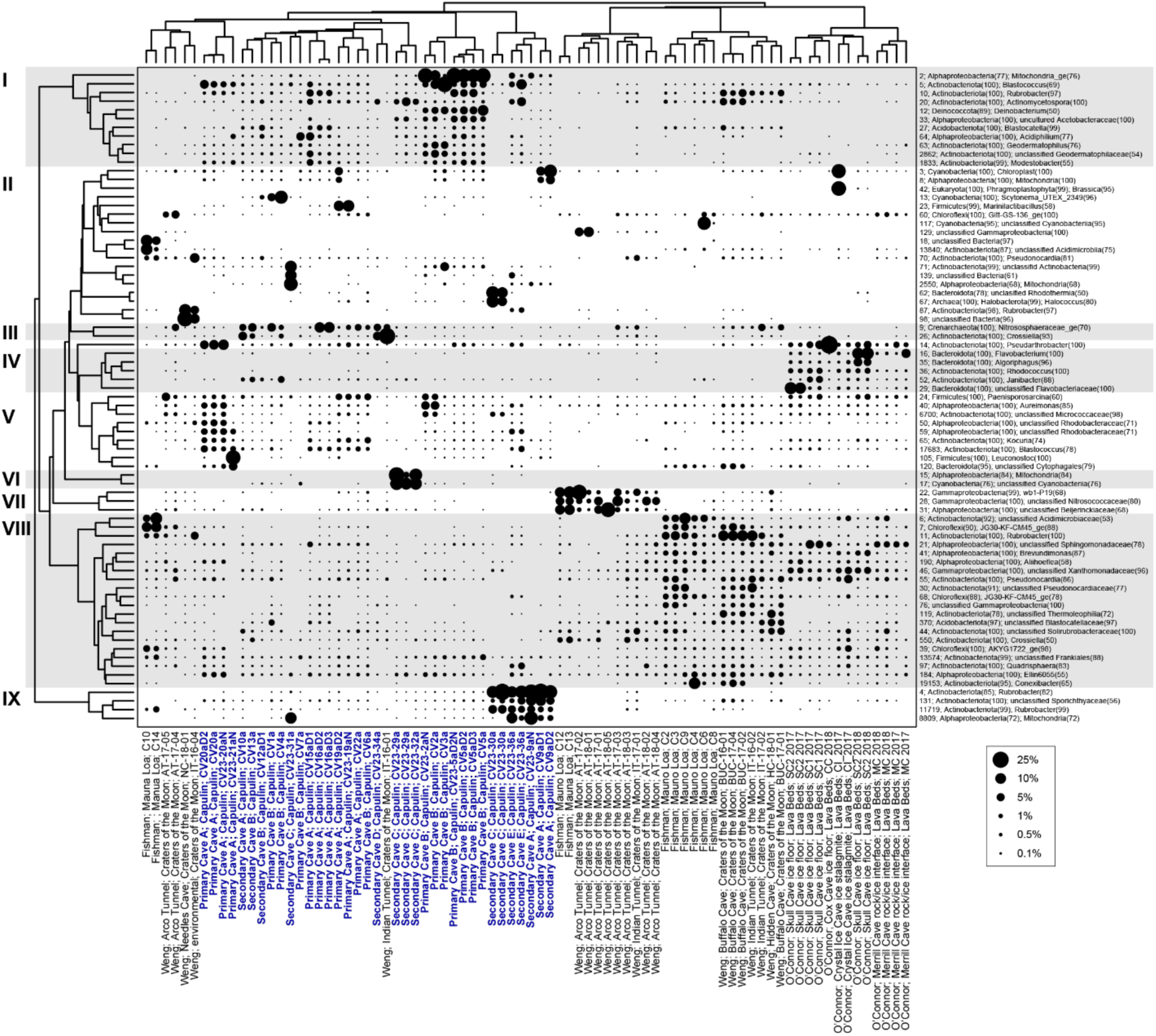
Two-way cluster analysis comparing rRNA gene libraries from Capulin Volcano to rRNA gene libraries from large lava tubes from three other studies: Mauna Loa, HI, USA (Fishman et al. 2023), Lava Beds National Monument, CA, USA (O’Connor et al. 2021), and Craters of the Moon, ID, USA (Weng et al. 2022). The R-mode analysis (clustering of taxa) was computed using the 75 most abundant OTUs, while the Q-mode analysis (clustering of libraries) was computed with all OTUs.

**Figure 7.**
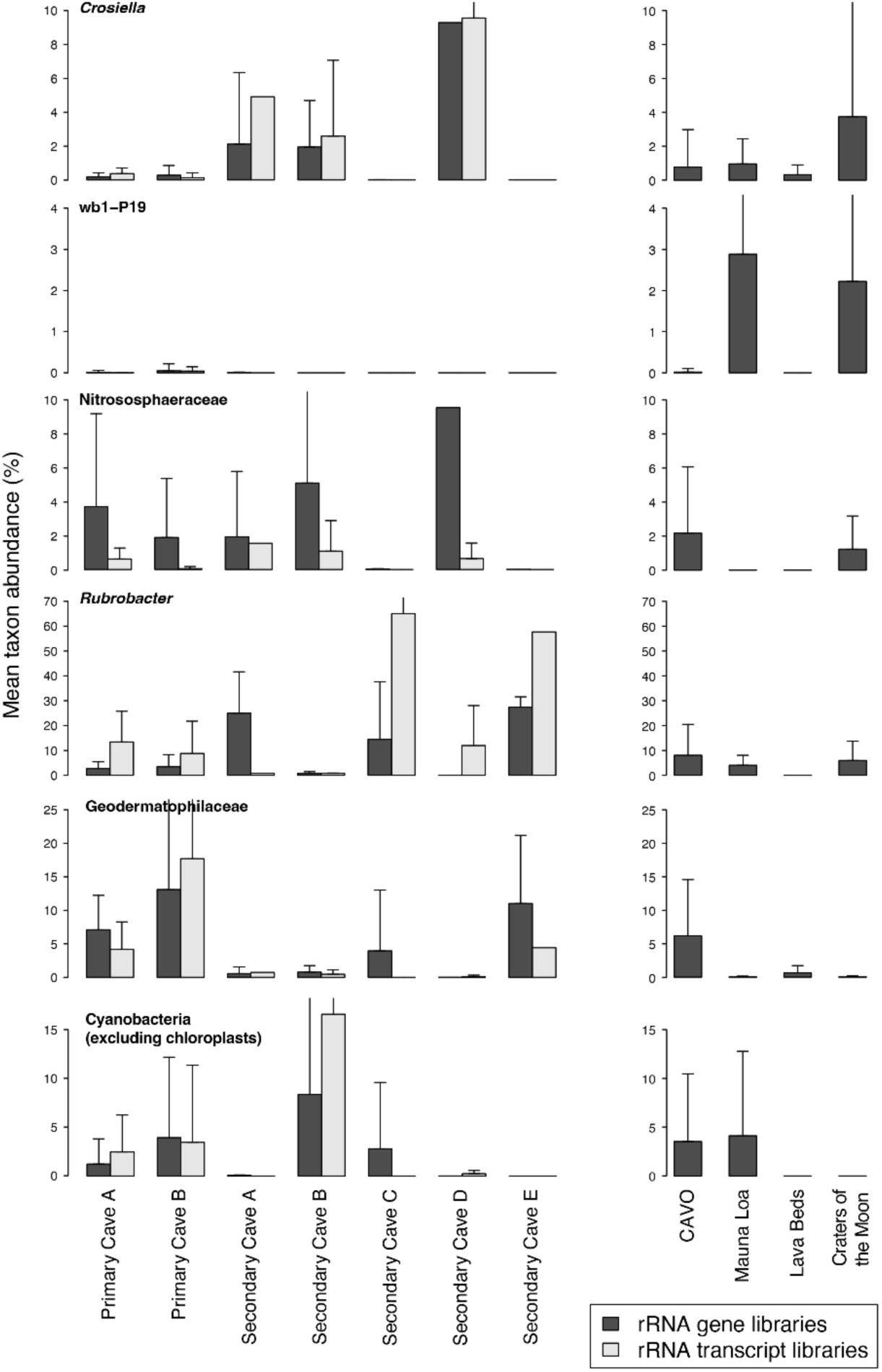
Mean abundance of select taxa across the difference CAVO caves, in both rRNA gene and transcript libraries. The panels to the right show the mean of each taxon across all libraries averaged from CAVO and the three other studies that were used for the comparison in Figure 5 and 6: Mauna Loa, HI, USA (Fishman et al. 2023), Lava Beds National Monument, CA, USA (O’Connor et al. 2021), and Craters of the Moon, ID, USA (Weng et al. 2022). Error bars show one standard deviation.

The most abundant microbial taxa include several abundant OTUs classified as *Rubrobacter* and *Modestobacter*. *Modestobacter* have been found in diverse environments, but are commonly associated with low water environments like Atacama Desert and Antarctic soils (Golińska et al. 2020), and many isolated *Rubrobacter* spp. are radiation and desiccation resistant (e.g., Ferreira et al. 1999; Egas et al. 2014). Several other abundant taxa are also often associated with harsh and arid environments such as *Deinobacter*, *Blastocatella*, *Kocuria*, and others (e.g., Ekman et al. 2011; Foesel et al. 2013; Xie et al. 2024; Cavanaugh et al. 2025). Most representatives of these abundant taxa have aerobic organoheterotrophic lifestyles, and we confirmed the presence of organoheterotrophs by obtaining several isolates R2A media (Table S2).

Although most of the abundant taxa are associated with organotrophic or phototrophic lifestyles, we successfully enriched microorganisms in a gradient tube culture designed for iron-oxidizing autotrophs, indicating that some chemolithotrophic populations are likely present. The presence of abundant OTUs classified as *Acidiphilium* (Figure 3) seems notable, as this is a genus often associated with metal- and sulfur-rich acid mine drainage environments, with some strains are capable of oxidizing sulfur compounds and reducing ferric iron (e.g., Johnson 2007; Li L et al. 2020). However, BLASTN analysis of the representative sequences of the most abundant OTUs shows that the closest isolates are strains AP34E and AP29E, which were isolated by Pulschen et al. (2017) from Antarctic soils using very dilute Diffco Nutrient Broth and long incubation times, so the *Acidiphilium*-like sequences identified here may represent a different lineage with ecological and physiological properties distinct from described *Acidiphilium* spp. Meta-omic or culture-based analyses will be necessary to further explore the metabolic capabilities of these lava cave-associated *Acidiphilium*-like taxa, as well as evaluate whether other populations are capable of lithoautotrophic growth.

### Microbial communities in the small lava caves at Capulin Volcano National Monument have many of the same taxa as larger lava tubes

Microbial communities in lava tubes have been much better characterized than those from small lava caves (e.g., Northup et al. 2011; Hathaway et al. 2014; Miller et al. 2014; Northup and Lavoie 2015; Gonzalez-Pimentel et al. 2021; O’Connor et al. 2021; Prescott et al. 2022; Weng et al. 2022; Nicolosi et al. 2023; Gutierrez-Patricio et al. 2024; Palma et al. 2024; Devlin et al. 2025; Kim Y et al. 2025; Salazar-Hamm et al. 2025). Smaller lava caves like those sampled here, the largest of which had less than 15 m of passage (Fig. 1), may not have the same environmental conditions as larger lava tubes. Lava tubes often extend kilometers in length with passages up to several meters tall that humans can walk through upright, and have dark zones, more stable temperatures, higher humidity, and different energy and nutrient resources than the surface (Culver and Pipan 2014; Northup and Lavoie 2015). Therefore, in order to determine if microbial communities from the small primary and secondary lava caves at CAVO reflect those of larger lava caves, we compared the 16S rRNA gene libraries from this study to those from three studies of lava tubes from different locations that used the V4 region. Because one of the studies we selected (O’Connor et al. 2021) used a different primer set than the others, and because the others had shorter read lengths, we only used forward reads for this comparison. It is possible that this difference in primers impacted the resulting comparison. In this analysis, libraries from O’Connor et al. (2021) do cluster separately from those of Weng et al. (2022) and Fishman et al. (2023) (Figure 6), although they are overall more similar to most libraries from these other lava tubes than to CAVO libraries (Figures 5 and 6), so it does not appear that primer biases substantially impact overall trends. We also note that we did not see a notable difference between CAVO libraries prepared using ‘full service’ versus ‘in house’ methods, which used slightly different primer sets, as most libraries prepared from the same sample cluster together (Figure 3).

Overall, microbial communities from the small CAVO caves have many of the same microbial populations as those from larger lava tubes, just with different proportions. The OTUs that occur in Clusters I, V, and IX in Figure 6 are more abundant in CAVO samples but still present in the lava tubes, and vice versa for the OTUs in Cluster VIII. If Cluster VIII represents microorganisms that are more likely to occur in cave-adapted assemblages, then perhaps the lower abundance of those taxa in CAVO caves reflect the greater surface influence of the smaller caves. For example, the genus *Crossiella* is thought to be an important group for biogeochemical carbon cycling in diverse cave systems (Gonzalez-Pimentel et al. 2021; Martin-Pozas et al. 2022; Martin-Pozas et al. 2023; Nicolosi et al. 2023; Gutierrez-Patricio et al. 2024), is also present in many CAVO libraries, but at relatively lower abundance (Figure 7). Other OTUs also follow this pattern. For example, *Brevundimonas* and “AKYG1722” occur in Cluster VIII, both of which are present but at lower abundance in CAVO libraries; Kim et al. (2025) identified *Brevundimonas* as generalists and “AKYG1722” as specialists in lava tube communities. In contrast, some of the most abundant genera in CAVO caves such as *Rubrobacter*, *Modestobacter* (a member of family Geodermatophilaceae), and unclassified members of the “Nitrososphaeraceae” are present but at lower abundance in the libraries from large lava tubes (Figures 6, 7).

Interestingly, while the gammaproteobacterial genus “wb1-P19” occurs in several samples from larger lava tubes, it was rare in CAVO libraries (Figure 7). wb1-P19 is commonly found in caves worldwide, and metagenomic analysis recently revealed that it is a methanotroph in the group “*Candidatus* Methyloligotrophales” that is likely existing on trace atmospheric methane (Jones et al. 2024; Bay et al. 2025). Perhaps increased light and organic carbon influence from the surface in small lava caves results in a more energy-rich environments where trace gas oxidizers like wb1-P19 are not competitive. Further comparisons of microbial communities among different cave zones and between large and small caves will be necessary to further determine cave-specific microbial assemblages worldwide.

### Implications for the geomicrobiology of lava caves

Although most studies on lava cave microbial ecology have focused on large lava tubes, small primary and secondary lava caves are common features in volcanic landscapes and may represent substantial subterranean void space in these environments. We found that many of the same groups of microorganisms found in large lava tubes also occur in small primary and secondary lava caves, indicating that these cave-associated populations may occupy more widespread habitats in basaltic lava flows, even if they are not as enriched as in large lava tubes. The presence of chloroplast sequences and other cyanobacteria in both large lava tubes and small lava caves (Figures 6, 7) suggests that light penetration, even if limited, supports phototrophic communities across a range of cave sizes. However, small lava caves appear to be dominated by aerobic organoheterotrophic taxa, with less evidence for taxa associated with lithotrophic metabolisms that are known from larger lava tubes (Selensky et al. 2021). Likewise, trace gas oxidizers such as wb1-P19 were not detected or had very low abundance in the small caves. Perhaps the environmental conditions in smaller voids do not favor these more specialized cave populations due to higher inputs of surface-derived organic carbon, light levels, or differences in air circulation. Future studies comparing the composition, activity, and metabolic potential of microbial communities across a broader range of cave and void sizes will help determine the factors that structure subsurface life in these settings and evaluate the collective contribution of small caves and voids to landscape-scale biogeochemical cycling in volcanic landscapes.

## Acknowledgements

Thanks to D. Kissner and the staff at Capulin Volcano National Monument for facilitating this research. B. Tobin provided preliminary observations of bat and invertebrate use in CAVO caves, A. Armstrong assisted with the light measurements in April 2026, and the staff at the University of Minnesota Genomics Center provided insightful discussion and assistance with amplicon sequencing of challenging samples. J. Hoberg, A. Baloun, J. Swift Jr., and A. Brown were supported by the Scientists-in-the-Parks (SIP) internship program during summer 2023. The SIP program is administered by National Park Service (NPS), in partnership with Stewards Individual Placements, the Ecological Society of America, and the Geological Society of America. Special thanks to C. Bitting and M. Hassey (formerly Ehmling) for coordination and assistance with the SIP program. Additional financial support was provided by the National Cave and Karst Research Institute (NCKRI), part of New Mexico Institute of Mining and Technology. Samples were collected under NPS permit CAVO-2023-SCI-0001.

